# Prevalent mouse phenotypes in the unexplored druggable genome

**DOI:** 10.1101/2021.11.08.467777

**Authors:** Olga Gulyaeva, Zicheng Hu, Tudor Oprea, K. C. Kent Lloyd, Shawn Gomez, Bryan L Roth, Michael T McManus

## Abstract

Among the estimated ~23,000 protein encoding human genes, the class of ‘druggable genes’– defined by their ability to bind drug-like compounds– represents an enticing collection of targets for clinical intervention. Yet many if not most of these genes remain poorly understood and understudied. Here we evaluate three major classes of druggable genes (GPCRs, ion channels, and kinases) and found that a third of these remain largely ignored yet display significant mouse phenotypes upon genetic ablation. We show that both well-studied and understudied druggable genes share a similar number and spectrum of phenotypes. Moreover, many of the mouse phenotypes arising from the ablation of both well-studied and understudied druggable genes show similarities with symptoms in rare human diseases. Collectively these data diminish the notion that most poorly studied genes may not be especially ‘important’ and highlight therapeutic opportunities and potential disease models among poorly characterized druggable genes.

## Introduction

After the first draft of human genome was published in 2000, the research community was presented a compendium of human genes, offering a veritable cornucopia of research opportunities for every annotated gene. However, in 2011 it appeared that approximately 75% of research activity still focused on only 10% of human genes, many of which were well-studied even before the human genome was published ^1,2^. It has been hypothesized that biomedical research is mostly guided by a small number of generic biochemical characteristics of genes rather than their physiological importance or relevance to human disease ^1–3^. It was also surmised that past research has a major influence on present research initiatives, e.g. 16% of human genes that had been known of prior to 1991 still accounted for ~50% of the literature in 2015 ^3^.

The mouse and human genome are each predicted to contain between 20-25,000 genes, of which at least ~3,000 genes have been previously classified as “druggable”, meaning that drug-like molecules may bind to their molecular nooks and crannies ^4^. Although druggability is difficult to define, this classification is largely based on gene families for which FDA-approved drugs already exist. Less than 25% of these druggable genes are currently targeted by FDA-approved drugs ^4,5^ further highlighting room for clinically relevant discoveries.

Within the druggable genome, GPCRs, ion channels and kinases particularly stand out as they represent the most successfully drug-targeted protein families based on the highest number of FDA-approved drugs developed against these genes ^6–8^. G-Protein-coupled receptors (GPCRs) represent a very diverse class of transmembrane integral proteins that serve a plethora of signaling functions in the cells ^8^. They typically act by interacting with a ligand which stabilizes particular receptor conformations, thereby recruiting other proteins that induce downstream signaling. Unlike other receptor classes, ligands for GPCRs are extremely diverse: they include ions, photons, vitamins, hormones, metabolites, and other proteins ^8,9^. GPCRs are known to induce their intracellular signaling upon ligand binding through indirect and direct interactions with ion channels, kinases and other proteins ^8^. There are over 400 genes encoding for non-olfactory GPCRs, many of which contribute to human diseases such as various neurological disorders and more recently discovered association to diabetes, obesity, heart disease and Alzheimer disease ^10^. It has been shown that over 30% of all drugs approved by the FDA act on GPCRs ^8^, highlighting their importance in disease and setting successful precedence for drug discovery.

Kinases represent another family of successfully drugged genes. Kinases are a class of enzymes or molecular “switches” that phosphorylate over 1/3 of the human proteome ^11,12^. There are close to 600 kinases encoded in the human genome and they have been implicated in many disease contexts including cancer, immunological, metabolic, cardiovascular and many other diseases. To date 76 kinase inhibitors have been approved by the FDA, with the majority targeting receptor tyrosine kinases for treatment of cancer ^13^. However, the vast majority of kinases lack these selective chemical probes, hence a large portion of this protein class remains poorly understood.

Ion channels regulate ion flux across cell- and organelle membranes and, thus, are involved in a plethora of cellular processes in the human body: membrane potential and excitability, neurotransmitter release, cell-to-cell interaction, cell volume and proliferation among other processes ^14,15^. Not surprisingly, this diverse class of proteins, comprised of >300 genes encoding various ion channel subunits ^16–18^ has been implicated in various diseases known as channelopathies. Over 60 channelopathies have been described to date, arising in a variety of human tissues such as the peripheral and central neural system, heart, skeletal muscle, kidney, bone and pancreas; these diseases may vary in the severity of their symptoms and in some cases can be lethal ^18^. It is worth noting, that although ~13.5% of the currently marketed drugs target ion channels, the majority of this class of genes remains poorly investigated ^19^.

By focusing on understudied targets similar to those for which drugs already exist (GPCRs, kinases, and ion channels), the NIH launched the Illuminating Druggable Genome (IDG) initiative in 2014. In this program the NIH defines “dark” by assessing the numbers of publications, grants and patents and in some cases by considering whether the “dark” protein has sufficient antibodies and drugs or modulators available for their study. In developing this program, the IDG initiative is attempting to address the persistent research biases among human genes, specifically– to promote emphasis on the study of genes that may have a more proximal translational impact through drug target discovery. In this program, the IDG hopes to identify potential new drug targets for medications to treat or cure some of the most burdensome diseases.

The IDG consortium has teamed with the Knockout Mouse Project (KOMP) and the International Mouse Phenotyping Consortium (IMPC). The KOMP uses newly available technologies and capabilities in high-throughput genome-wide mutagenesis to create knockout (conditional and deletion) alleles for every human gene with a mouse ortholog in mouse embryonic stem (ES) cells ^20^. This led to an international effort known as the International Mouse Phenotyping Consortium (IMPC) to use these and other partner ES cell collections to functionally annotate the entire mammalian protein coding genome with an emphasis on genes for which little annotation exists including the druggable genes. The IMPC produces male and female cohorts of gene knockout mice using ES cells and more recently CRISPR/Cas9 technologies applied directly in embryos for broad, non-targeted “discovery” phenotyping involving ~500 tests and procedures at 21 research laboratories in 15 countries on 5 continents around the globe. All mice generated are deposited in publicly accessible biorepositories and data is freely available and accessible at https://www.impc.org. To date, knockout mice for over 9,000 genes have been produced with more than 7,500 phenotyped thus far. Most mice show measurable phenotypes, many with extensive pleiotropy and sexual dimorphism. Approximately one third of the genes are either embryonic lethal or subviable. Yet with the tens of thousands of genes and an immeasurable number of potential phenotyping assays, the research community is only ‘scratching the surface’ in the understanding of gene function. And although the translational relevance to human biology is often intact, the ability to intercede with therapeutic intervention for human disease remains a distant vision.

Perhaps there is a biological basis for the gene study bias in the human genome. It has been suggested that well-studied genes might display more pronounced phenotypes in various biological studies and organismal models. According to this hypothesis, genes displaying the most obvious phenotypes might have been studied first, implying that most poorly studied genes may not be especially ‘important’ ^21–23^. Here we examine IMPC phenotypic datasets to quantitate and compare the number and nature of phenotypes between dark druggable genes and well-studied druggable genes. We show evidence that both dark and well-studied genes share a similar number and spectrum of phenotypes, divorcing the general idea that poorly studied genes may just not be that ‘important’. We illuminate novel mouse phenotypes for all three major classes of druggable genes highlighting the importance of studying dark druggable genes and encouraging researchers to consider KOMP and IMPC resources to better understand biological roles for these human disease models.

## Results

The human genome comprises 23,266 protein coding genes, out of which 8750 could be druggable based on predicted protein function and if similar genes in the family have been targeted by an FDA-approved drug ^24^ (Figure S1a, left diagram and Supplementary Table 1). Approximately 16% (1390) of these druggable genes belong to non-olfactory GPCRs, ion channels and kinases – the most successfully druggable gene classes to date. In its 2020 revision, the IDG settled on a final list of understudied (“dark”) human genes from these 3 classes, consisting of 152 GPCR, 102 ion channel and 162 kinase genes (Fig S1a, right diagram). Since mouse orthologues do not exist for every single of these human genes, the total number of mouse genes in each class is slightly lower: 117, 91 and 121 for GPCRs, ion channels and kinases respectively. As expected, on average dark genes have lower number of antibodies, patents, publications and ligands associated with each gene compared to those that are better studied (i.e., “light”) genes in these three classes (Figure S1b, top).

To evaluate whether dark druggable genes contribute to mouse phenotypes similar to light druggable genes, the number of knockout mouse phenotypes per each of the GPCRs, ion channels and kinases was analyzed and compared with Jensen score associated with each gene (Fig S1b bottom). The Jensen score quantifies the fractional count of protein mentions derived from text mining PubMed abstracts ^25^. On average, knockouts of genes with both low and high Jensen score associated with significant number of phenotypes, indicating that mouse phenotypes are not only caused by ablation of genes with higher Jensen score and therefore higher degree of characterization. We further examined how many mice were produced and phenotyped to date for dark and light genes. For light genes, mice were produced and phenotyped for 186 GPCRs out of the whole class of 304 light GPCRs, for 120 out of the class of 237 of light ion channels and for 281 out of the class of 502 of light kinases (Figure S1c left plots in each category). Thus, for over one third of the annotated genes in each light gene class, knockout mice were produced and phenotyped. Similarly, for over a third of dark genes, mice were produced and phenotyped: for 64 out of 117, 42 out of 91, 49 out of 121 for dark GPCRs, ion channels and kinases respectively (Figure S1c right plots in each category), albeit with a lower total number of genes in the dark list. These data show no obvious bias in preferentially phenotyping mice resulting from genetic ablation of better studied versus dark genes, likely reflecting efforts of IDG consortium to guide KOMP in prioritization to generate and phenotype KO mice for dark genes in addition to better studied ones.

The numbers of significant phenotypes associated with each gene were next analyzed. On average, genetic knockout of light genes in each class resulted in slightly higher rates of significant phenotypes than dark genes: 2.23 vs 1.85, 2.41 vs 1.53, and 3.13 vs 3.00 phenotype for light vs dark genes in GPCR, ion channel and kinase classes respectively (Figure S1d). This difference was not statistically significant and can be partially explained by higher average number of tests performed on KO mice for better studied genes compared to dark genes (Figure S1e). Perhaps this phenomenon can be attributed to inherent bias, where researchers tend to phenotype and characterize mice harboring deletions of genes for which there is already solid biological evidence. However, once significant phenotypes were expressed as a percentage among examined phenotypes, there were no significant differences between dark and light genes (Figure S1f).

Mouse phenotypes were subjectively binned into twelve categories, defined by tissue or organ system (Fig 1). Among these categories, a comparison of the well-studied and dark genes gave no indication that ablation of better studied genes causes more phenotypes than dark genes. The compendium of knockout mouse phenotypes for GPCRs, ion channels and kinases revealed a wide spectrum of affected tissues for both dark and well-studied genes. Cluster analysis for both dark and light genes based on phenotypic similarity demonstrated that both light and dark genes contribute significantly to a plethora of phenotypes with no apparent bias towards light genes (Fig 2). These data show that both dark and light genes similarly contribute to mouse phenotypes.

**Figure 1.**
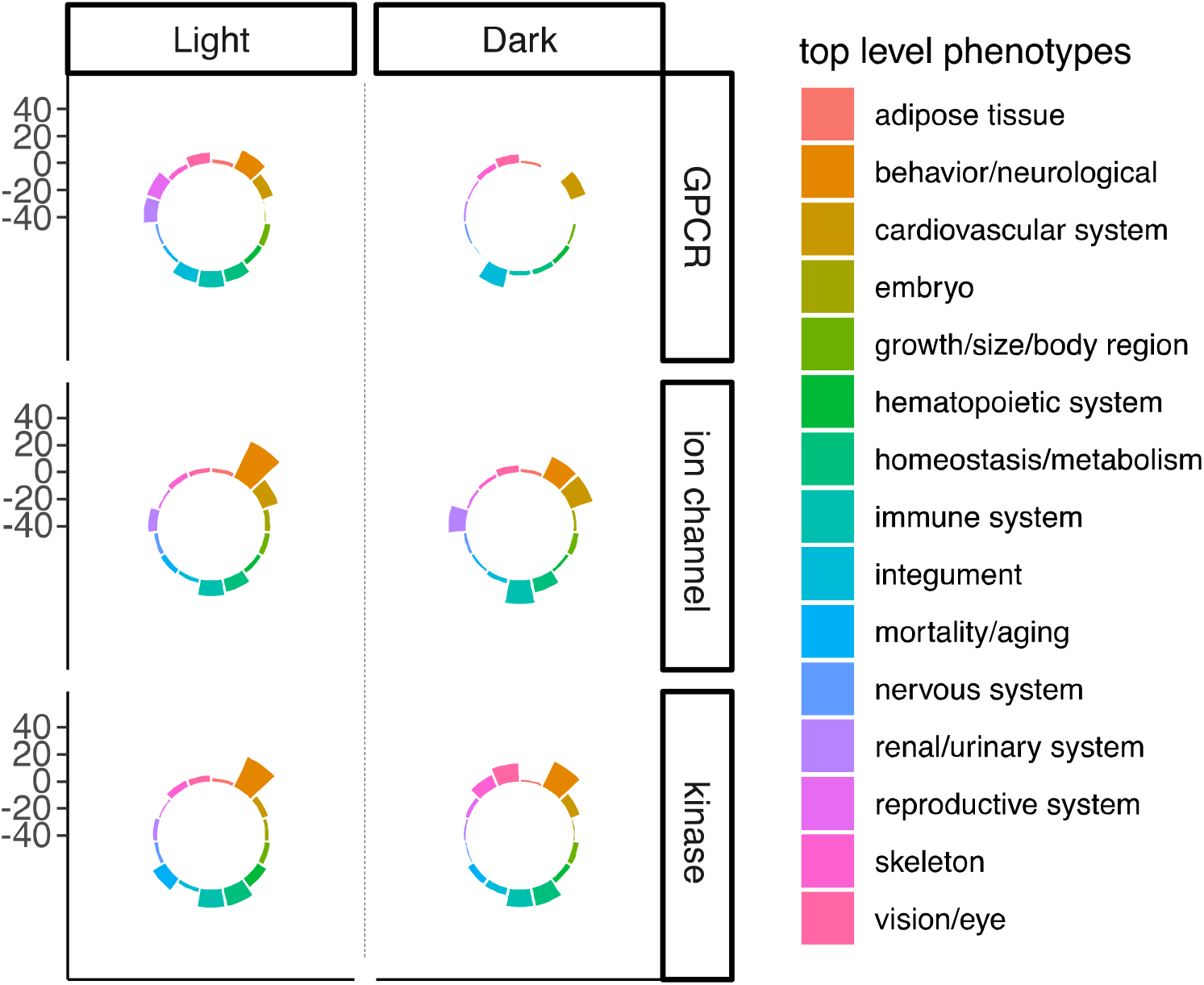
Quantification of significant mouse phenotypes for light and dark genes. Circular plots show the percent of significant mouse phenotypes per each dark and light gene class grouped by phenotypic category.

**Figure 2.**
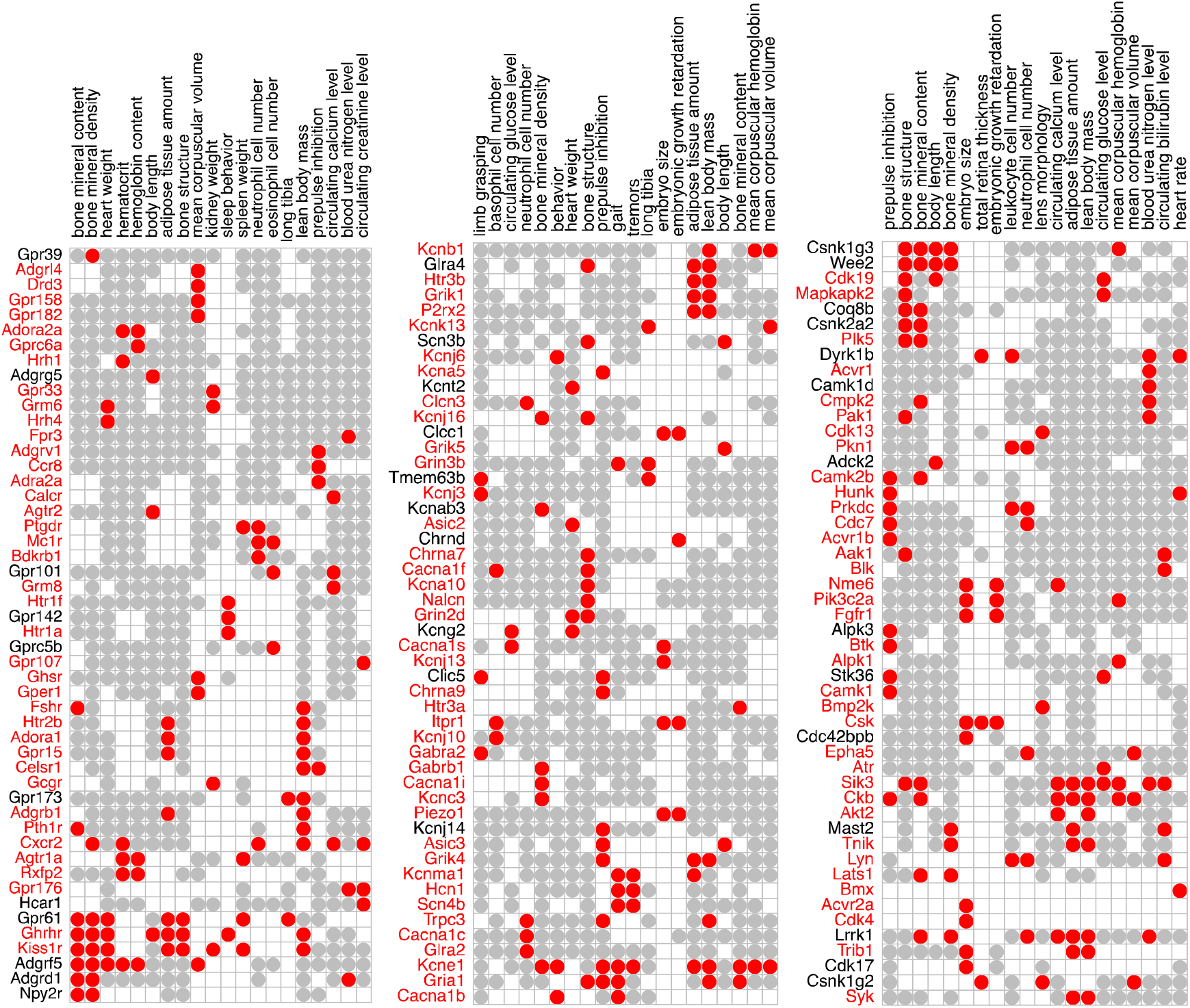
Examples of significant phenotypes for selected dark and light genes. For visualization, the top 20 most frequently measured phenotypes and the top 50 most thoroughly phenotyped knockout mice are shown in the heatmap. The heatmaps on the left, center and right show GPCR, ion channels and kinases, respectively. Red represents significant phenotype, grey represents non-significant phenotype, and absence of color indicates phenotype that has not been examined by the IMPC. Dark genes are depicted in black and light genes are depicted in red.

To determine whether well-studied genes have stronger human disease associations than dark genes, ClinVar, OMIM and GWAS catalog were mined. These data showed that ~30-60% of dark genes were significantly associated with either a monogenic or a polygenic disease: e.g., 37/126 (29%), 52/102 (51%), 76/125 (61%) of GPCRs, ion channels and kinases respectively (Supplementary Table 2 and Figure 3 top). A similar overall percentage of light genes showed disease association: 162/282 (57%), 162/245 (66%) and 357/510 (70%) for GPCRs, ion channels and kinases respectively. These data suggest that although many dark genes are relevant for human health, they have been historically ignored likely due to the lack of prior knowledge, interest, or availability of resources to interrogate these genes.

**Figure 3.**
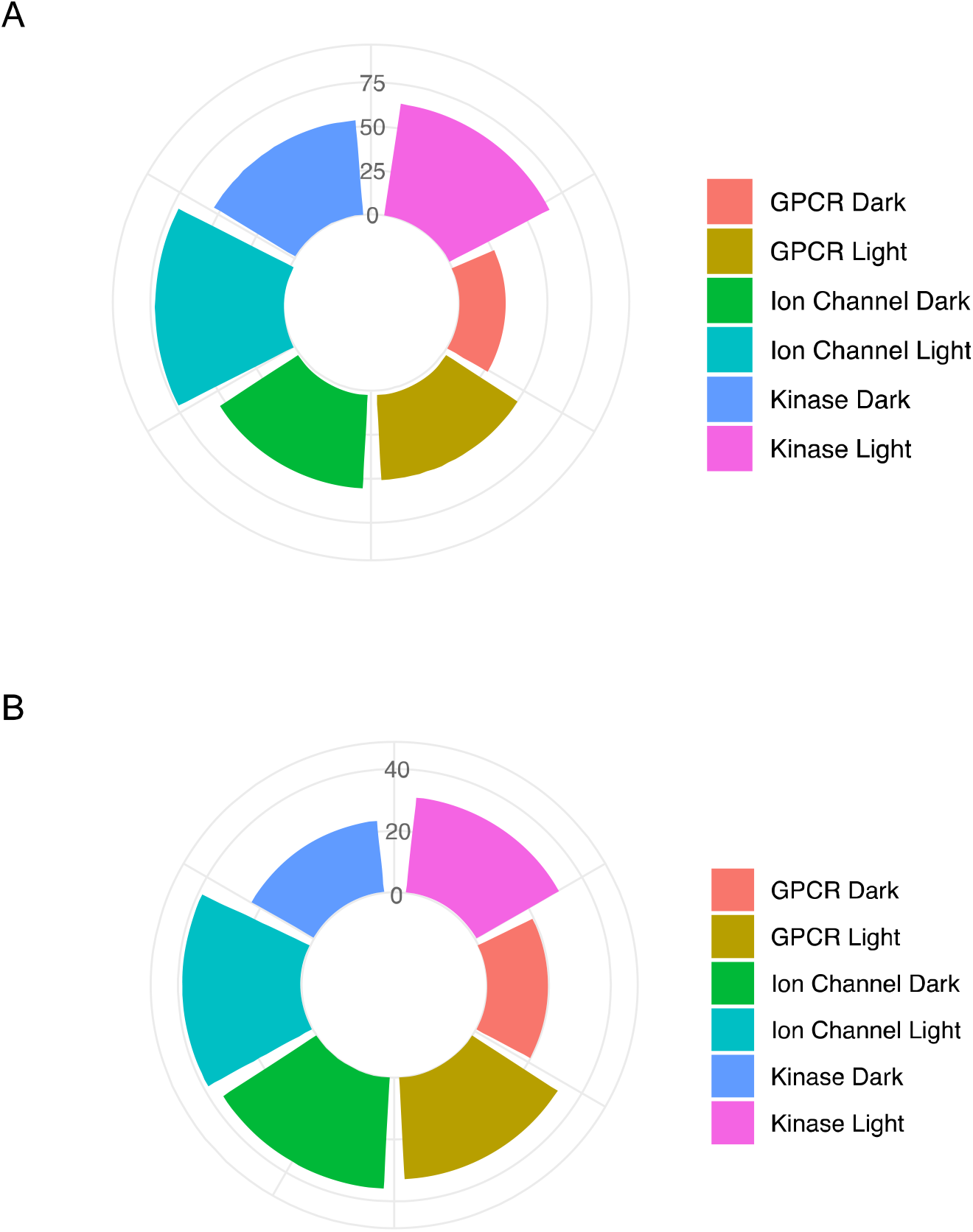
Association of druggable genes with human diseases. (A) Bar graphs depicting percentage of genes in each class associated with human diseases. (B)Bar graph depicting percentage of genes for which a mouse model serves as a reliable model for human disease based on Phenodigm score > 90.

To predict which gene knockout mouse models might serve as models for specific human diseases, knockout mouse phenotypes were compared with human disease symptoms and a similarity score between the two was calculated using a previously developed method “Phenodigm” ^26^. This analysis identified a significant number of potential disease models for both dark and light genes (Fig 3 bottom and Supplementary Table 3), pointing to potential therapeutic relevance for dark genes. Dark genes with the most significant mouse phenotypes upon ablation matching to similar phenotype and disease in humans were highlighted in the context of a particular tissue in the body (Figure 4, Figure S1g and Supplementary Table 4). To assess how well dark genes can tolerate loss of function mutation in the human population, a comparison to light genes was performed using NOMAD data. Dark genes contained loss-of-function mutations at a similar frequency when compared to well-studied light genes (Figure S1h). Overall, these results demonstrate that mouse models are extremely informative for both light and dark genes and can provide a solid model system to study human diseases and aid in development of novel therapeutics.

**Figure 4.**
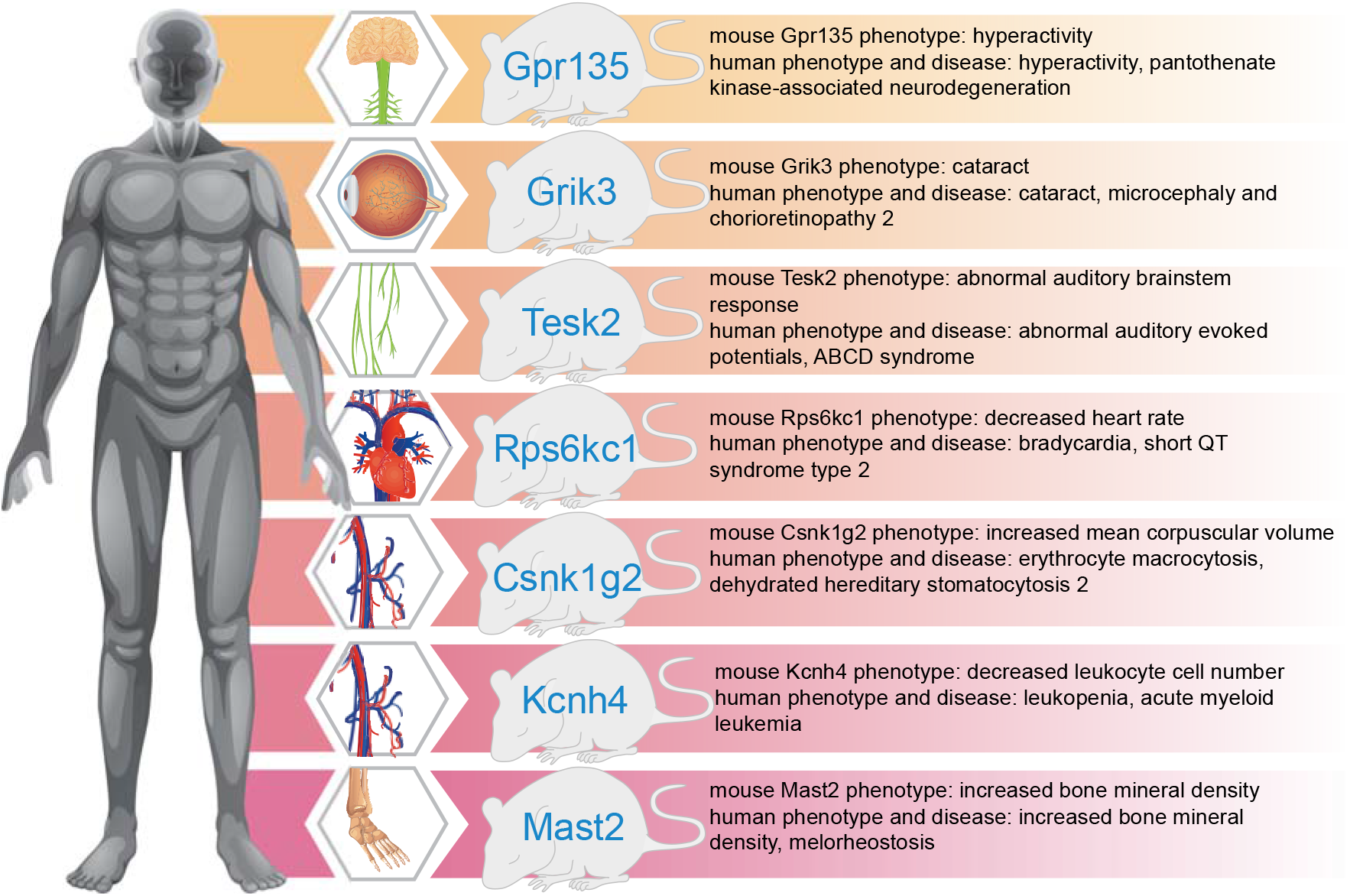
Summary of the most significant dark genes from IMPC data set for which similar phenotypes were found in humans. Phenotypes associated with each human disease were identified using human phenotype ontology (HPO) and significant mouse phenotypes were identified using high-throughput phenotype data from the International Mouse phenotype consortium (IMPC). Mouse phenotypes were matched with human phenotypes using the mappings provided by the PhenomeNET (Hoehndorf et al., 2011).

## Discussion

Historically, researchers have turned to model organisms to interrogate gene function ^27,28^. Here the aim was to emphasize novel and important mouse phenotypes arising from dark gene ablation in combined efforts of IDG and KOMP. Dark druggable genes from 3 classes studied here (GPCRs, ion channel and kinases) resulted in many significant phenotypes upon ablation in mice proving their importance in various cellular processes. Most of these dark genes had very little known about their function prior to the IDG launch in 2014: in many cases there were few or no peer reviewed publications or grants about them thus yielding in relatively low Jensen score. Indeed, many significant phenotypes observed could not have been predicted based on protein sequence or expression pattern available through public databases.

With the help of IMPC/KOMP phenotyping data, genes from IDG dark list can now be prioritized for their in-depth study in a particular cellular and organ context. Additionally, genes with significant phenotypes in mice and/or disease association in humans represent best therapeutic targets for diseases. For example, IDG dark list of 102 dark ion channels was parsed out into “high-priority” list consisting of 15 genes for more focused cellular and *in vivo* studies largely based on the following criteria: a) significant mouse phenotypes as identified by KOMP; b) human disease association; c) unexpected cellular properties caused by ion channel overexpression (unpublished data). When compiling this list, particular attention was paid to genes that are very dark and whenever possible, family members were also included to complete the gene class. Full list of prioritized dark genes for cellular and mouse studies can be found in Supplementary Table 6. This allows to focus scientific effort on a subset of genes for certain biological questions for which these genes are likely the most suitable or relevant.

Similarly, some genes from IDG dark list may be deprioritized if there was no significant phenotype observed for that particular gene KO. However, it is worth noting that absence of phenotypes does not mean that the gene is not important or functional in other contexts/cells/tissues that simply were not covered by phenotyping tests performed by KOMP. Moreover, for some gene classes with high sequence similarity among class members, such as ion channels and kinases, the lack of phenotype may arise due to compensation from a homologous gene. In addition, the interconnected nature of kinase signaling can lead to no observable phenotype due to the activation of compensatory signaling pathways upon kinase knockout. This is a highly practical concern as such plasticity is a major component of drug resistance and reprogramming in cancer ^29,30^. Additionally, some genes may not show phenotype in the mouse strain that KOMP used to generate all their mice (C57Bl6), while they may have significant phenotypes in other strains. In fact, some of the genes described in this manuscript may have other phenotypes described by individual investigators that may not be reflected in the IMPC data which could be caused by different mouse backgrounds, statistical analysis methods, and the fact that IMPC does not examine all possible phenotypes. But despite these caveats, IMPC/KOMP added significant new and previously unknown or unexpected functional annotations to dark genes and tremendously helped IDG centers to prioritize genes for study using various assays. Finally, all IDG and KOMP data are freely available online, and mice and other resources are available for a fee from relevant repositories; all mouse strains are cryopreserved and stored in MMRRC [https://www.mmrrc.org], all IDG resources are stored in public repositories [https://darkmatter.ucsf.edu/ and https://pharos.nih.gov/] or available directly from labs upon request, see also Supplementary Table 5. All KOMP data is available at https://www.impc.org.

Overall, we found that dark and well-studied druggable genes share a similar number and spectrum of mouse phenotypes upon genetic ablation and are similarly functionally conserved, thus dismissing the general notion that dark genes may not be as informative for human health. We also show that mouse models are extremely informative for both light and dark genes and can provide a solid model system to study human diseases and aid in development of novel therapeutics. Most of the conventional drugs (small molecule and antibodies) possess inhibitory action on proteins they target. However, most genetic diseases are caused by loss-of-function mutations, therefore it is desirable to activate rather than inactivate these genes. Thus, research programs centered around generating activating drugs for druggable genome targets are needed and would aid significantly in treatment of channelopathies as well as other diseases caused by GPCRs and kinases.

## Supporting information

Supplemental informaton

## Acknowledgements

We thank Lily Jan, John Gilchrist, all members of the McManus Lab who provided critical feedback during the course of this project. This project was supported by NIH IDG fund 1U24 DK116214-01.

## Contributions

The project was designed and conceived by OG, MTM, and KL. OG wrote the manuscript. ZH performed all bioinformatic analysis and data mining TIO provided critical feedback and analyses. All authors read and approved the final manuscript.

## Methods

### Data sources

The mouse phenotype data is downloaded from the International Mouse Phenotyping Consortium (https://www.mousephenotype.org) ^31^, data release 14 was used for Figure 1 and S1c-f and S1h, while data release 13 was used for Figure 2 and S1g and Supplementary Table 3. To identify mouse models for each human disease, we queried the phenodigm API (https://www.mousephenotype.org/help/programmatic-data-access/phenodigm-results), which provides the similarity score between the phenotype of a knock-out mouse and the symptoms of a human disease ^26^. The Jensen scores of the druggable genes are downloaded from the Pharos portal (https://pharos.nih.gov) ^32^. Human diseases associations with druggable genes were mined in ClinVar, OMIM and GWAS catalogs (https://www.ncbi.nlm.nih.gov/clinvar/, https://www.omim.org/, https://www.gwascentral.org/). Information related to druggable protein families are downloaded from the Drug Target Ontology (http://drugtargetontology.org/) ^24^. The frequency of the predicted loss-of-function mutations for each gene is downloaded from the genomAD (https://gnomad.broadinstitute.org/) ^33^.

### Statistical analysis

A p-value threshold of 0.0001 is used to identify significant mouse phenotypes from mouse phenotype dataset according to the recommendation from the IMPC consortium. Non-parametric unpaired two-samples Wilcoxon test is used to test the quantitative differences (number of phenotypes, number of disease models and frequencies of loss-of-function mutations) between dark and non-dark genes. Bonferroni corrections are used to adjust for multiple comparisons.

### KOMP

Methods for producing and phenotyping genes are harmonized using agreed standard operating procedures and protocols across the three KOMP2 Centers. Briefly, knockout mice were generated from gene-targeted ES cells or by CRISPR/Cas9 genome editing directly in zygotes using standard procedures. After gene confirmation and additional quality control procedures, cohorts of 7-8 male and female homozygous (or heterozygous in the case of embryonic lethal genes) mice were produced for phenotyping beginning at 4 weeks of age, including weekly body weights, fertility and viability testing, and a set of multiple observations. Cohorts were analyzed for abnormalities in 11 body systems between 9 and 15 weeks of age after which mice were euthanized for gross necropsy, histopathology, and blood analysis. A subgroup of genes was selected at random for aging and repeated phenotyping again between 49 and 70 weeks of age to identify late adult-onset abnormalities. Embryo lethal and subviable genes were studied at various gestational ages of development using gross, microscopic specialized imaging (uCT) technologies to identify the approximate time and morphologic evidence for the cause of death. The study was conducted in accordance with the guidelines set forth by the 8th Edition of the Guide for the Use and Care of Laboratory Animals and the Public Health Service Policy on Humane Care and Use of Laboratory Animals, and was approved by the University of California Davis Institutional Animal Care and Use Committee under the Protocol for Animal Care and Use #20863.

### Data and Code availability

The mouse phenotype data are available on the International Mouse Phenotyping Consortium website (https://www.mousephenotype.org). All codes used to analyze data in the current study are available from the corresponding author upon request.

## Citations

1. Edwards, A. M. et al. Too many roads not taken. Nature 470, 163–165 (2011).

2. Oprea, T. I. et al. Unexplored therapeutic opportunities in the human genome. Nat. Rev. Drug Discov. 17, 317–332 (2018).

3. Stoeger, T., Gerlach, M., Morimoto, R. I. & Amaral, L. A. N. Large-scale investigation of the reasons why potentially important genes are ignored. PLOS Biol. 16, e2006643 (2018).

4. Hopkins, A. L. & Groom, C. R. The druggable genome. Nat. Rev. Drug Discov. 1, 727 (2002).

5. Rodgers, G. et al. Glimmers in illuminating the druggable genome. Nat. Rev. Drug Discov. 17, 301–302 (2018).

6. Overington, J. P., Al-Lazikani, B. & Hopkins, A. L. How many drug targets are there? Nat. Rev. Drug Discov. 5, 993–996 (2006).

7. Santos, R. et al. A comprehensive map of molecular drug targets. Nat. Rev. Drug Discov. 16, 19–34 (2017).

8. Wacker, D., Stevens, R. C. & Roth, B. L. How Ligands Illuminate GPCR Molecular Pharmacology. Cell 170, 414–427 (2017).

9. Roth, B. L. Molecular pharmacology of metabotropic receptors targeted by neuropsychiatric drugs. Nat. Struct. Mol. Biol. 26, 535–544 (2019).

10. Hauser, A. S., Gloriam, D. E., Bräuner◻Osborne, H. & Foster, S. R. Novel approaches leading towards peptide GPCR de-orphanisation. Br. J. Pharmacol. 177, 961–968 (2020).

11. McClendon, C. L., Kornev, A. P., Gilson, M. K. & Taylor, S. S. Dynamic architecture of a protein kinase. Proc. Natl. Acad. Sci. 111, E4623–E4631 (2014).

12. Ferguson, F. M. & Gray, N. S. Kinase inhibitors: the road ahead. Nat. Rev. Drug Discov. 17, 353–377 (2018).

13. Cohen, P., Cross, D. & Jänne, P. A. Kinase drug discovery 20 years after imatinib: progress and future directions. Nat. Rev. Drug Discov. 20, 551–569 (2021).

14. Isacoff, E. Y., Jan, L. Y. & Minor, D. L. Conduits of life’s spark: A perspective on ion channel research since the birth of Neuron. Neuron 80, (2013).

15. Subramanyam, P. & Colecraft, H. M. Ion Channel Engineering: Perspectives and Strategies. J. Mol. Biol. 427, 190–204 (2015).

16. Jegla, T. J., Zmasek, C. M., Batalov, S. & Nayak, S. K. Evolution of the human ion channel set. Comb. Chem. High Throughput Screen. 12, 2–23 (2009).

17. Bagal, S. K. et al. Ion Channels as Therapeutic Targets: A Drug Discovery Perspective. J. Med. Chem. 56, 593–624 (2013).

18. Imbrici, P. et al. Therapeutic Approaches to Genetic Ion Channelopathies and Perspectives in Drug Discovery. Front. Pharmacol. 7, (2016).

19. Garcia, M. L. & Kaczorowski, G. J. Ion channels find a pathway for therapeutic success. Proc. Natl. Acad. Sci. 113, 5472–5474 (2016).

20. Brown, S. D. M. & Moore, M. W. The International Mouse Phenotyping Consortium: past and future perspectives on mouse phenotyping. Mamm. Genome 23, 632–640 (2012).

21. Hoffmann, R. & Valencia, A. Life cycles of successful genes. Trends Genet. 19, 79–81 (2003).

22. Pfeiffer, T. & Hoffmann, R. Temporal patterns of genes in scientific publications. Proc. Natl. Acad. Sci. 104, 12052–12056 (2007).

23. Haynes, W. A., Tomczak, A. & Khatri, P. Gene annotation bias impedes biomedical research. Sci. Rep. 8, 1362 (2018).

24. Lin, Y. et al. Drug target ontology to classify and integrate drug discovery data. J. Biomed. Semant. 8, 50 (2017).

25. Pletscher-Frankild, S., Pallejà, A., Tsafou, K., Binder, J. X. & Jensen, L. J. DISEASES: Text mining and data integration of disease–gene associations. Methods 74, 83–89 (2015).

26. Smedley, D. et al. PhenoDigm: analyzing curated annotations to associate animal models with human diseases. Database J. Biol. Databases Curation 2013, bat025 (2013).

27. Hunter, P. The paradox of model organisms. The use of model organisms in research will continue despite their shortcomings. EMBO Rep. 9, 717–720 (2008).

28. Goldstein, B. & King, N. The Future of Cell Biology: Emerging Model Organisms. Trends Cell Biol. 26, 818–824 (2016).

29. Gross, S., Rahal, R., Stransky, N., Lengauer, C. & Hoeflich, K. P. Targeting cancer with kinase inhibitors. J. Clin. Invest. 125, 1780–1789 (2015).

30. Manstein, V. von & Groner, B. Tumor cell resistance against targeted therapeutics: the density of cultured glioma tumor cells enhances Stat3 activity and offers protection against the tyrosine kinase inhibitor canertinib. MedChemComm 8, 96–102 (2017).

31. Koscielny, G. et al. The International Mouse Phenotyping Consortium Web Portal, a unified point of access for knockout mice and related phenotyping data. Nucleic Acids Res. 42, D802–809 (2014).

32. Sheils, T. et al. How to Illuminate the Druggable Genome Using Pharos. Curr. Protoc. Bioinforma. 69, e92 (2020).

33. Karczewski, K. J. et al. The mutational constraint spectrum quantified from variation in 141,456 humans. Nature 581, 434–443 (2020).

