## Supplemental informaton for "Prevalent mouse phenotypes in the unexplored druggable genome"

**Supplementary Figure 1. Dark druggable genes represent a prevalent part of human genome and contribute to mouse knockout phenotypes.** (attached separately)

A. Proportion of dark druggable gene classes in the human genome. B. Top: Average number of antibodies, patents, publications and ligands associated with each gene in dark and light class. Bottom: Number of mouse phenotypes caused by gene ablation plotted against the genes’ Jensen score. This graph shows no apparent correlation between Jensen score (and therefore darkness of the gene) with number of phenotypes associated with that gene. Therefore, dark genes, although remain understudied, contribute to mouse phenotypes. Raw numbers used to draw these plots can be found in Supplementary Table 1. C. Number of KO mice phenotyped in each gene class. D. Number of significant KO mouse phenotypes observed in each gene class. E. Number of total KO mouse phenotypes examined in each gene class. F. Percent of significant KO phenotypes among examined phenotypes. G. Examples of raw phenotypic data associated with each gene from Figure 1b right. H. Violin plots depicting percentage of observed loss of function mutations (LOF) in light or dark gene classes and their relationship to mouse phenotypes according to gnomAD database. The Y-axis shows the precent individuals predicted to carry loss-of-function mutations in the population based on gnomAD dataset (Karczewski et al., 2020). The color of the dots represents if a significant (p value <0.0001) phenotype has been identified by the high-throughput mouse phenotype pipeline of the IMPC.

**Supplementary Tables**

**Supplementary Table 1. List of genes in the human genome classified into druggable and dark categories.**

| **family** | **number** | **percentage** |
| --- | --- | --- |
| druggable | 8750 | 37.60853 |
| others | 14516 | 62.39147 |
| **family** | **number** | **percentage** |
| Surfactant | 5 | 0.05714286 |
| Storage | 8 | 0.09142857 |
| Nuclear receptor | 48 | 0.54857143 |
| Cell-cell junction | 61 | 0.69714286 |
| Extracellular structure | 65 | 0.74285714 |
| Chaperone | 68 | 0.77714286 |
| Cell adhesion | 68 | 0.77714286 |
| Epigenetic regulator | 91 | 1.04 |
| Other families | 414 |  |
| Calcium-binding protein | 117 | 1.33714286 |
| Immune response | 123 | 1.40571429 |
| Signaling | 427 | 4.88 |
| Cellular structure | 437 | 4.99428571 |
| Transcription factor | 760 | 8.68571429 |
| Nucleic acid binding | 789 | 9.01714286 |
| Transporter | 890 | 10.17142857 |
| Receptor | 326 | 3.72571429 |
| Ion channel | 347 | 4.02285714 |
| GPCR | 408 | 4.64 |
| Kinase | 635 | 6.60571429 |
| Enzyme modulator | 705 | 8.05714286 |
| Other Enzymes | 2426 | 27.72571429 |
| **family** | **number** | **percentage** |
| Light GPCR | 282 | 20.273185 |
| Dark GPCR | 126 | 9.058231 |
| Light ion channel | 245 | 17.613228 |
| Dark ion channel | 102 | 7.332854 |
| Light kinase | 510 | 36.66427 |
| Dark kinase | 125 | 8.986341 |

**Supplementary Table 2. List of human disease associations for dark genes (attached separately).**

**Supplementary Table 3. List of mouse models that serve as human disease models.**

| **gene** | **mouse_homolog** | **idgFamily** | **diseaseTerm** | **score** |
| --- | --- | --- | --- | --- |
| GPR173 | Gpr173 | GPCR | Acroleukopathy, Symmetric | 90.4 |
| GPRC5C | Gprc5c | GPCR | Anemia, Sideroblastic, 2, Pyridoxine-Refractory | 90.595 |
| GPR39 | Gpr39 | GPCR | Polycystic Kidney Disease 5 | 92.54 |
| GPR174 | Gpr174 | GPCR | Autoimmune Disease | 92.58 |
| GPR19 | Gpr19 | GPCR | Lithium Transport | 93.54 |
| LPAR6 | Lpar6 | GPCR | Thalassemia, Beta+, Silent Allele | 93.93 |
| NPY2R | Npy2r | GPCR | Bulimia Nervosa, Susceptibility To | 96.08 |
| TAS2R14 | Tas2r116 | GPCR | Cystic Angiomatosis Of Bone, Diffuse | 100 |
| MRGPRX4 | Mrgprb2 | GPCR | Cataract 29 | 100 |
| GPR68 | Gpr68 | GPCR | Danubian Endemic Familial Nephropathy | 100 |
| TAS2R13 | Tas2r102 | GPCR | Adrenocortical Carcinoma, Hereditary | 100 |
| KCNT1 | Kcnt1 | Ion channel | Pruritus, Hereditary Localized | 90.45 |
| CACNG6 | Cacng6 | Ion channel | Ribbing Disease | 92.215 |
| CATSPER2 | Catsper2 | Ion channel | Spermatogenic Failure 5 | 93.145 |
| SCN3B | Scn3b | Ion channel | Cardiac Conduction Defect | 93.27 |
| FXYD3 | Fxyd3 | Ion channel | Blood Group--Lutheran Inhibitor | 93.54 |
| KCNMB4 | Kcnmb4 | Ion channel | Electroencephalogram, Low-Voltage | 96.08 |
| SCN2B | Scn2b | Ion channel | Maturity-Onset Diabetes Of The Young, Type 13 | 96.165 |
| CACNB2 | Cacnb2 | Ion channel | Maturity-Onset Diabetes Of The Young, Type 13 | 96.165 |
| KCNV1 | Kcnv1 | Ion channel | Iris Pigment Epithelium Anomalies | 96.61 |
| TMC7 | Tmc7 | Ion channel | Oocyte Maturation Defect 2 | 98.215 |
| CACNG3 | Cacng3 | Ion channel | Hypercholesterolemia, Familial, 3 | 98.61 |
| SCNN1B | Scnn1b | Ion channel | Developmental Dysplasia Of The Hip 1 | 100 |
| TMEM38B | Tmem38b | Ion channel | Genitourinary Tract Anomalies | 100 |
| KCNJ15 | Kcnj15 | Ion channel | Charcot-Marie-Tooth Disease, Axonal, Type 2V | 100 |
| KCNAB3 | Kcnab3 | Ion channel | Ribbing Disease | 100 |
| PKD2L2 | Pkd2l2 | Ion channel | Attention Deficit-Hyperactivity Disorder | 100 |
| TMEM63A | Tmem63a | Ion channel | Benign Hereditary Chorea | 100 |
| GLRB | Glrb | Ion channel | Nanophthalmos 2 | 100 |
| PAK6 | Pak6 | Kinase | Acroleukopathy, Symmetric | 90.4 |
| NEK10 | Nek10 | Kinase | Acroleukopathy, Symmetric | 90.4 |
| MAP3K14 | Map3k14 | Kinase | X-Linked Lymphoproliferative Disease | 90.9 |
| RPS6KL1 | Rps6kl1 | Kinase | Anemia, Sideroblastic, Pyridoxine-Responsive, Autosomal Recessive | 91.285 |
| PIP5K1B | Pip5k1b | Kinase | Nondisjunction | 91.38 |
| PHKG2 | Phkg2 | Kinase | Erythrocytosis, Familial, 4 | 91.735 |
| SCYL1 | Scyl1 | Kinase | Neuropathy, Hereditary Sensory, Atypical | 91.785 |
| CDK14 | Cdk14 | Kinase | Hypomagnesemia 4, Renal | 92.24 |
| PSKH1 | Pskh1 | Kinase | Maturity-Onset Diabetes Of The Young, Type 13 | 96.165 |
| TSSK6 | Tssk6 | Kinase | Spermatogenic Failure 2 | 97.255 |
| CSNK2A2 | Csnk2a2 | Kinase | Male Infertility Due To Acephalic Spermatozoa | 97.515 |
| STK33 | Stk33 | Kinase | Spermatogenic Failure 5 | 98.435 |
| HIPK4 | Hipk4 | Kinase | Spermatogenic Failure 5 | 98.435 |
| PRKCQ | Prkcq | Kinase | Platelet Aggregation, Spontaneous | 98.735 |
| CDKL2 | Cdkl2 | Kinase | Cephalin Lipidosis | 100 |
| MAST4 | Mast4 | Kinase | Malocclusion Due To Protuberant Upper Front Teeth | 100 |
| SCYL3 | Scyl3 | Kinase | Exercise Intolerance, Riboflavin-Responsive | 100 |
| NEK5 | Nek5 | Kinase | Tune Deafness | 100 |

**Supplementary Table 4. List of genes with matching phenotypes between mice and humans (attached separately).**

**Supplementary Table 5. List of all IDG associated websites.**

| Types | Website name | Description | URL |
| --- | --- | --- | --- |
| IDG generated resources and data | Protein illumination timeline | The data portal for resources and data generated by the three IDG Data and Resource Generation Centers, including GPCR, kinase, ion channel groups. | https://druggablegenome.net/ProteinTimeLine |
|  | Dark matter | The data portal for resources and data generated by the IDG ion channel Data and Resource Generation Center. | https://darkmatter.ucsf.edu/ |
|  | A Mouse Imaging Server (AMIS) | An imaging server to browse the mouse imaging data generated by IDG GPCR Data and Resource Generation Center. | https://amis.docking.org/ |
|  | International mouse phenotype consortium | A data portal for mouse phenotypes profiled by the International Mouse phenotype Consortium. | https://www.mousephenotype.org/ |
| Informatics tools | Pharos | Pharos is the user interface to the Knowledge Management Center (KMC) for the Illuminating the Druggable Genome (IDG) program. | https://pharos.nih.gov/ |
|  | GENEVA | GENEVA (GENe Expression Variance Analysis) is a semi-automated framework for exploring public RNA-seq datasets. GENEVA allows researchers to identify RNA-seq datasets that contain modulating conditions for a gene or a gene signature. | https://genevatool.org/ |
|  | Reactome | Pathway-based analysis and visualization of understudied human proteins. | https://reactome.org/ |
|  | Monarch initiative | The Monarch Initiative is an integrative data and analytic platform connecting phenotypes to genotypes across species, bridging basic and applied research with semantics-based analysis. | https://monarchinitiative.org/ |
|  | Drug central | DrugCentral provides information on active ingredients, chemical entities, pharmaceutical products, drug mode of action, indications, pharmacologic action. | http://drugcentral.org |
|  | Dark kinome knowledgebase | Dark Kinome Knowledgebase generates, systematizes and disseminates knowledge about dark kinases, biological networks in which they function and connections to cellular phenotypes and human disease. | https://darkkinome.org/ |
|  | Protein kinase ontology browser | Protein Kinase Ontology Browser allows users to locate protein kinase proteins and a lot of information related to the proteins, including the sequence, structure, function, mutation and pathway information on kinases. | http://vulcan.cs.uga.edu/prokino/about/prokino |

**Supplementary Table 6. List of prioritized dark ion channel genes and their characteristics (attached separately).**
