## Supplementary figures and images for "Prevalent mouse phenotypes in the unexplored druggable genome"

### Supplemental Figure 1

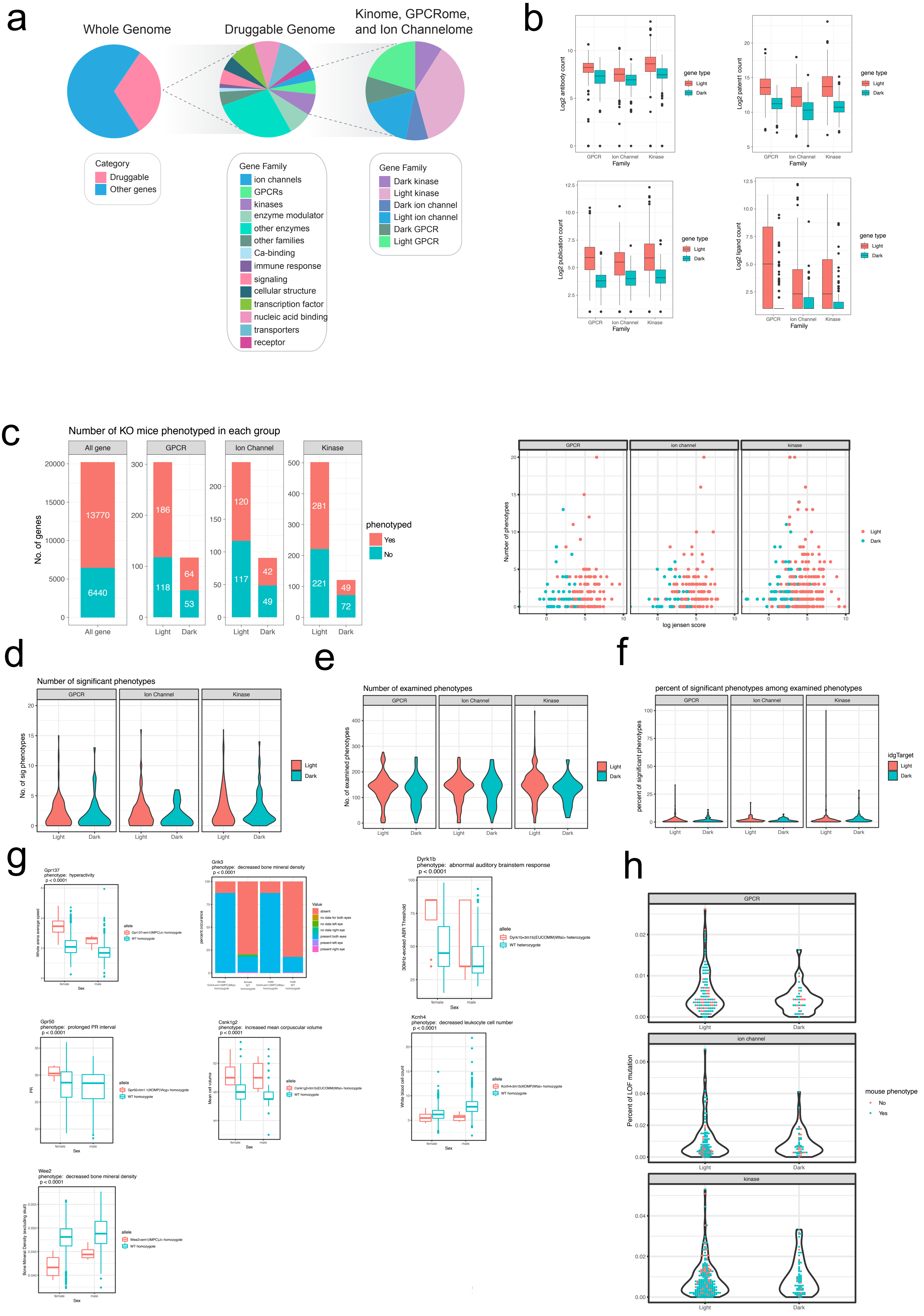
